# Epidermal threads reveal the origin of hagfish slime

**DOI:** 10.1101/2022.07.06.499062

**Authors:** Yu Zeng, David Plachetzki, Kristen Nieders, Hannah Campbell, Marissa Cartee, Kennedy Guillen, Douglas Fudge

## Abstract

Fiber-reinforced soft materials possess high flexibility with high strength but are rare in nature. Hagfishes can produce a tough, fibrous slime within a fraction of a second by ejecting two cellular products, mucus and threads, into seawater. With thousands of silk-like threads, the slime is highly effective in defending against large predators. However, the evolutionary origin of hagfish slime remains unresolved, with the presence of another, putatively homologous thread in the epidermis providing circumstantial evidence for an epidermal origin. Here, we investigated the epidermal threads produced in hagfish skin. We found that these threads average ∼2 mm in length and ∼0.5 μm in diameter, or ∼80 times shorter and ∼4 times thinner than the slime threads, characterizing the second longest intracellular fiber. The entire hagfish body is covered by a dense layer of epidermal thread cells, with each square millimeter of skin storing a total of ∼96 cm threads. Experimentally induced damage to a hagfish’s skin caused the release of threads, which together with mucus, formed an adhesive epidermal slime that is more fibrous and less dilute than the defensive slime. Transcriptome analyses further revealed that the epidermal threads are ancestral to the slime threads, with duplication and diversification of thread genes in parallel with the evolution of slime glands. These results support an epidermal origin of hagfish slime and slime glands, as driven by predator selection for stronger and more voluminous slime.

## 1. Introduction

Composite materials are produced from two or more constituent materials for improved, novel performance. Fiber-reinforced soft materials possess special properties (e.g., high viscoelasticity and flexibility with high strength) and have broad applications, such as medical biomaterials and tissue engineering (Pan et al., 2007; Gil et al., 2013; O’Brien, 2011). However, there are very few fibrous soft materials found in nature, partly due to the physical challenge of effective mixing (e.g., the requirement of turbulence for mixing particles and fibers with fluid; Hishida et al., 1992; Olson, 2001).

Among the various defensive structures used by animals, hagfish slime is a fibrous hydrogel, recognized by its exceptional material properties and unique deployment mechanisms (Ewoldt et al. 2011; Chaudhary et al. 2018; Lim et al. 2006; Winegard et al. 2010, Bernards et al. 2018). The hagfishes (Class Myxini) are a group of jawless vertebrates inhabiting the ocean floor as scavengers and predators. A row of slime glands occurs along each side of a hagfish, with each gland producing and storing gland thread cells (GTCs) and gland mucous cells (GMCs). When a hagfish is attacked, it produces defensive slime by rapidly ejecting ruptured GMCs and GTCs into seawater (**Fig. 1A**). Within 400 ms after ejection, coiled threads from GTCs unravel and mucous vesicles from GMCs swell and deform, resulting in a network of mucus and threads that entraps large volumes of water and effectively clogs the mouth and gills of fish predators (Lim et al. 2006; Zintzen et al. 2011).

**Figure 1.**
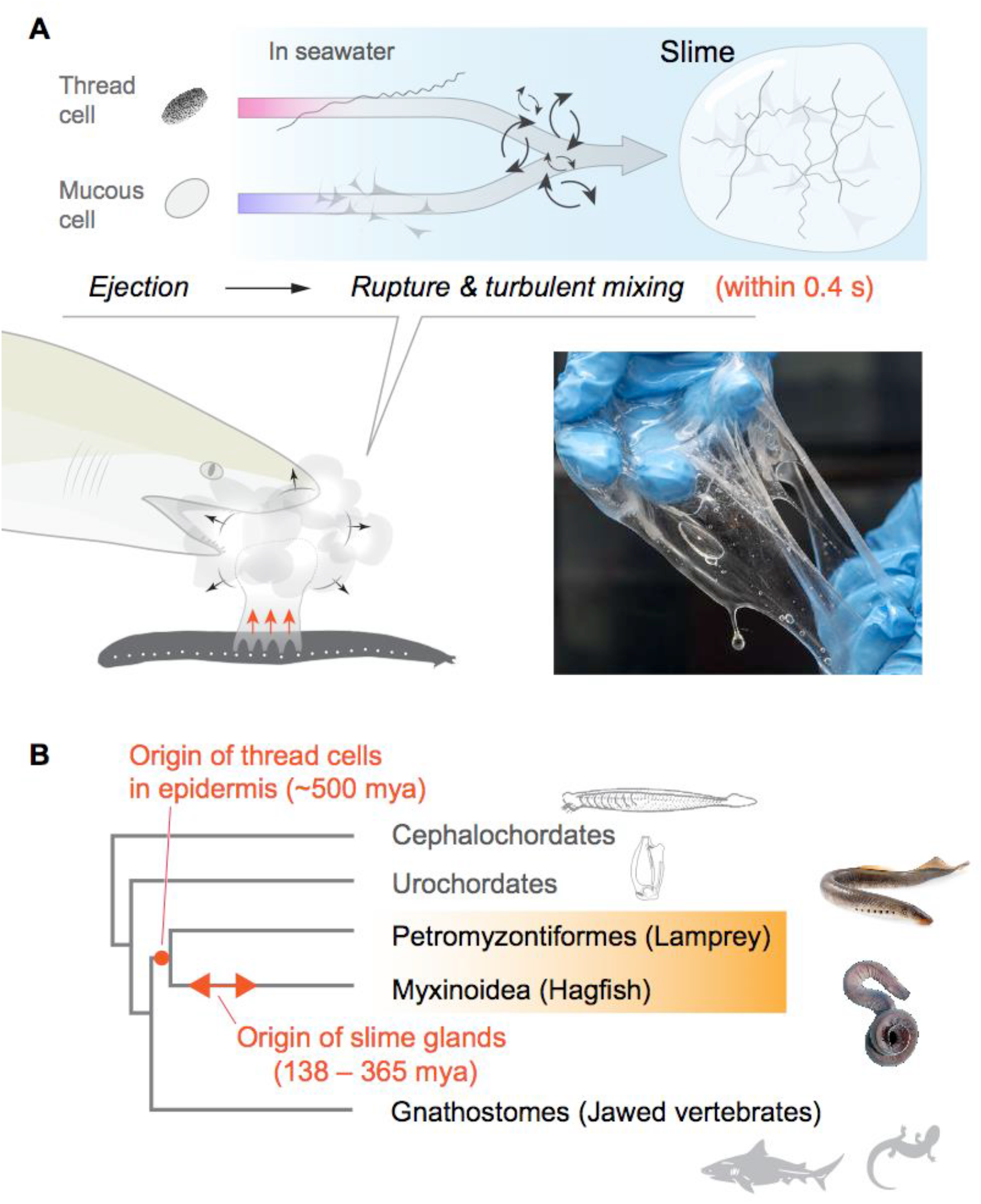
Mechanism and evolutionary history of hagfish slime. **(A)** Hagfish defensive slime is produced by rapid ejection and rupture of mucous cells and thread cells into seawater by slime glands. Top shows a schematic sequence of slime formation. Threads and mucus are released from ruptured cells and mix with seawater to form large volumes of dilute, soft, viscoelastic slime (lower right). **(B)** A simplified cladogram of chordates annotated with the origin of hagfish slime gland and the presence of epidermal thread cells (orange shade). Fossil evidence suggests hagfish slime glands evolved between 138 – 365 million years ago (mya; Miyashita, 2020). Thread-producing epidermal cells are found in only hagfishes and lampreys. This suggests a single origin of epidermal thread cells in their common ancestor (Cyclostomi), which dated to ∼500 mya and likely preceded the hagfish slime glands.

In spite of being composed mostly of water (i.e., > 99.99% seawater), hagfish slime is strong and viscoelastic (Fudge et al., 2003; Fudge et al., 2005; Ewoldt et al. 2011; Fudge et al., 2015; Böni et al., 2016). A single pinch on the tail of an adult Pacific hagfish (*Eptatretus stoutii*; ∼45 cm body length) can cause the production of 0.9 liters of slime, which is ∼7 times the volume of the animal (∼0.14 liters) (Fudge et al., 2005). There is no other biological or synthetic that can expand so much in so little time, and there has been no artificial fibrous material that is as dilute and strong as hagfish slime. The effectiveness of defensive slime may have helped the hagfishes persist through the rise and dominance of the jawed fishes, while most other jawless vertebrates have gone extinct (Randle and Sansom, 2019).

The impressive strength of the slime is imparted by a network of slime threads that, together with webs of mucus, entrain large volumes of seawater. The threads are proteinaceous fibers, individually produced and stored within GTCs as a densely packed skein (Downing et al., 1981a,b; Winegard et al., 2014). Slime threads consist mainly of fibrous α and γ proteins from the intermediate filament family, and they rival spider silk in their strength and toughness (Downing et al., 1984; Spitzer et al., 1988; Koch et al., 1995; Fudge et al., 2003; Fudge and Gosline, 2004; Fudge et al. 2010). A recent study showed that larger hagfishes produce longer and thicker slime threads, presumably to defend against larger predators. With diameter and length varying four-fold (0.7 – 4 μm and 5 – 22 cm, respectively), hagfish slime threads are the largest intracellular polymers known in biology (Zeng et al., 2021).

The fossil record for hagfishes is sparse, which makes tracing the evolutionary origins of hagfish slime glands difficult. Available fossil data suggest that slime glands appeared in hagfishes somewhere between 138 – 365 million years ago (Miyashita, 2020). While the presence of thread-producing cells in the epidermis of both lampreys and hagfishes suggests an origin of threads in epidermis preceding the origin of hagfish slime glands (**Fig. 1B**).

Anatomical studies of extant hagfishes also suggest that slime glands arose as modifications and internalizations of the epidermis. An unusual thread-producing cells in hagfish epidermis - epidermal thread cells (ETCs) – were suspected to be homologous to GTCs in slime glands. Early studies showed that each ETC produces a single thread loosely packed within the cytoplasm (Schreiner, 1916). Ultrastructural studies revealed that immature ETC threads consist of a bundle of filaments that are 8 – 14 nm in diameter and resemble cytoplasmic intermediate filaments (Blackstad, 1963). Moreover, unlike GTCs, ETCs produce a dense mass of granules of unknown function in the distal region of the cell (Schreiner, 1916). While ETC threads and granules appear to be secretory products, there is no evidence that ETCs secrete threads or granules via merocrine or apocrine modes (Blackstad, 1963).

If the threads produced by ETCs are destined for export, a mechanism that depends on cell rupture is more likely, and this would be consistent with the holocrine secretion of threads and mucus that occurs in the slime glands. There is, however, no obvious mechanism of ETC rupture and release, such as the muscle fibers that surround the slime glands.

We thus hypothesized that ETCs rupture and release their contents when the skin is damaged, especially during interactions with predators. Hagfish skin is flaccid and allows them to survive bites from sharp-toothed predators such as sharks (Boggett et al., 2017). Although the deployment of slime can effectively deter these predators, its release is only triggered after the initial attack. Thus, hagfishes are likely to sustain frequent damage to their skin from predator bites under natural conditions (Zintzen et al., 2011), and the rupture of epidermal cells may resemble the release of alarm cues during skin damage in lampreys and many jawed fishes (Pfeiffer and Pletcher, 1964; Bals and Wagner, 2012; Pandey et al., 2021).

To address the function of epidermal threads, we collected morphological and experimental data from hagfish skin. We first quantified the abundance and morphology of epidermal threads and then examined the slime product on hagfish skin after experimentally inducing damages. We found the ruptured epidermal thread cells released threads and granules, forming an adhesive slime that was more fibrous and concentrated than the defensive slime. With transcriptomics analysis, we found the thread biopolymers in hagfish skin are ancestral to those found in slime glands, with gene duplication and divergence generating a diversity of thread biopolymers uniquely expressed in slime glands. With clear evidence suggesting an epidermal origin of hagfish slime, we further derived a general model to explain the initial evolution of hagfish slime glands as driven by predator selection.

## 2. Results

### 2.1 Thread and mucous cells cover the entire hagfish body

Within an epidermal thickness of approximately 95 – 110 μm, epidermal thread cells (ETCs) are generally found in the basal half (∼50 μm and deeper) of the epidermis, along with large mucus cells (LMCs). ETCs and LMCs are covered by 3 – 5 layers of small mucus cells (SMCs) (**Fig. 2A**). Viewing the skin perpendicular to the apical surface (*en face* view), the outer epidermal surface is covered by densely packed SMCs, while the deeper portion contains mainly ETCs and LMCs (**Movie S1**,**S2**). To assess the abundance of ETCs over the entire epidermis, we sampled the density of all three epidermal cell types from nine transverse cross-sections from head to tail (**SI Fig. S1C-E**). We approximated the area density of each cell type with respect to skin area as *σ* = *λ*^2^, where *λ* is the linear density sampled from the cross-section of skin.

**Figure 2.**
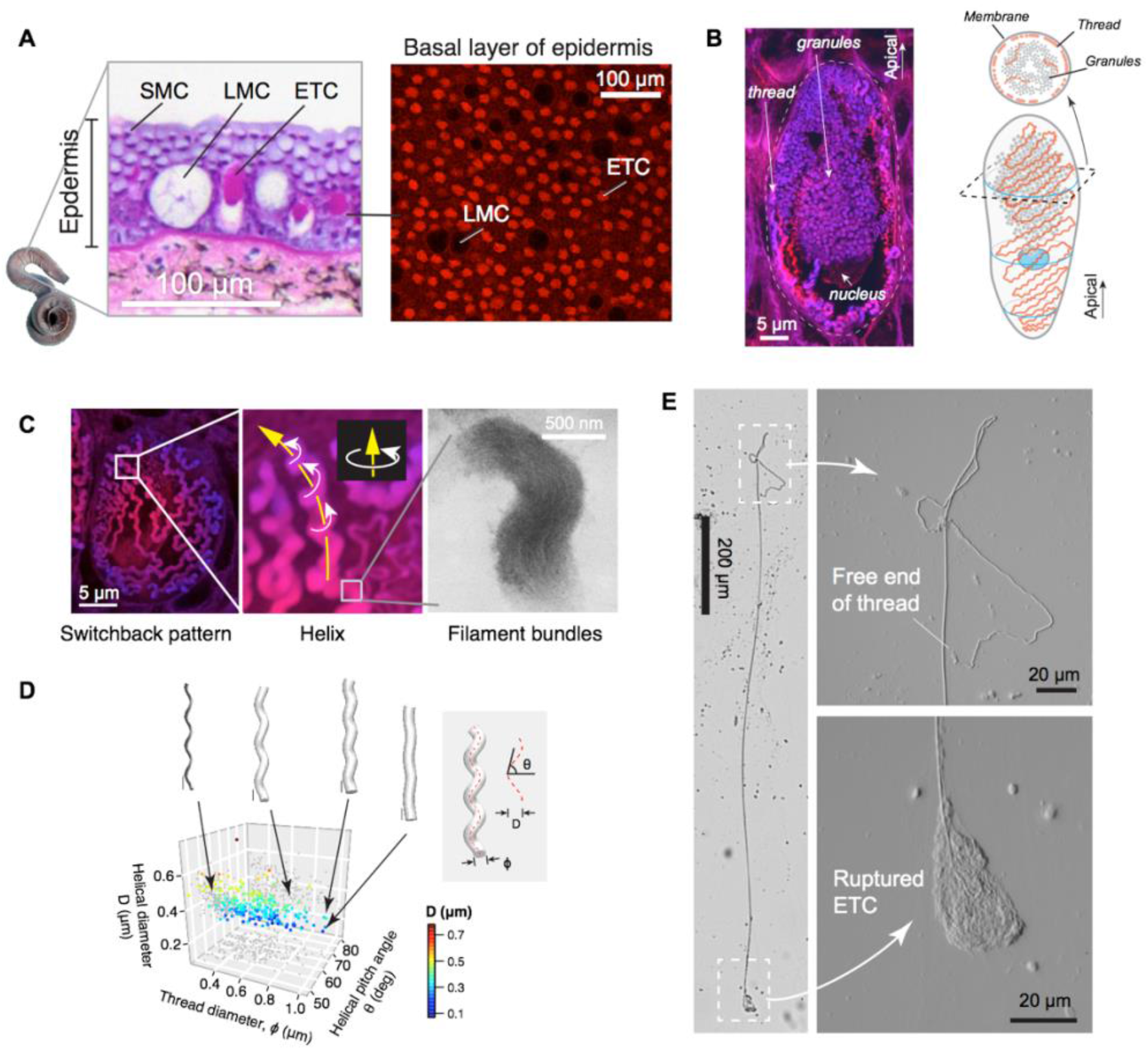
Hagfish epidermal threads. **(A)** Cross-section of dorsal epidermis from a Pacific Hagfish (*Eptatretus stoutii*; hematoxylin-eosin stained). (Right) The basal layer of epidermis containing epidermal thread cells (ETCs) and large mucus cells (LMCs), as captured with confocal microscopy in *en face* view. ETCs are characterized by granules and threads stained with the fluorescent stain eosin; LMCs appear as circular voids. **(B)** Longitudinal cross-section of an ETC, showing a cluster of granules, the nucleus located at the basal region of the granules, and a helical thread located mainly along the inner surface of the plasma membrane. (Right) Schematic of major cellular components of an ETC. **(C)** Three levels of epidermal thread structure. (Left - Middle) At the micro-scale, the thread traces a right-handed helix, the centerline of which is arranged in a switchback pattern on the inner surface of the cell membrane. Yellow arrow denotes the direction of increase; white arrows denote direction of helical rotation. (Right) At the nano-scale, a thread consists of a dense bundle of intermediate filament proteins, shown here in TEM (see also **SI Fig. S3D**). **(D)** Variations in thread geometry with respect to a morpho-space defined by thread diameter *ϕ*, helical pitch angle *θ* and helical diameter *D*. With increasing pitch angle *θ*, thread diameter *Φ* increases (*P* < 0.05; linear regression model) and helical diameter *D* decreases (*P* < 0.001), illustrated with idealized threads (scale bars, 1 μm). **(E)** A partially released thread (∼2 mm long) from a ruptured ETC, as viewed under light microscopy (see also **SI Fig. S3A**).

We found that the proportion of the three cell types varies little across the different regions sampled, with ETCs being the second most abundant. The mean area density of ETCs was ∼434 mm^-2^. For an adult hagfish (∼45 cm long), we estimate a total of ∼ 1.2 × 10^7^ ETCs covering the entire hagfish body. Notably, this total number of ETCs is ∼3.9 times greater than the total number of GTCs from all slime glands combined (∼3.1 × 10^6^, assuming a total of 163 glands; see **SI Appendix A**). In addition, the LMCs occurred with density ∼92 mm^-2^, which is ∼4.7 times lower than that of ETCs. The SMCs occurred at a density of 4.3 × 10^5^ mm^-2^, which is ∼1000 times more abundant than the ETCs. (**SI Fig. S1E**,**F**). These abundance data allowed us to approximate the relative proportions of cellular products in epidermal mucus (see below).

### 2.2 Structure of epidermal thread cells

We also examined cross-sections of *E. stoutii* skin using laser scanning confocal microscopy and identified three prominent structures within ETCs: (1) a densely packed granule cluster, (2) a thread that is loosely packed along the inner plasma membrane and also interweaves among granules, and (3) a large nucleus located at the basal surface of the granule cluster (**Fig. 2B**). Such a layout features more unoccupied cytoplasmic space compared to GTCs, which are mostly occupied by the nucleus and thread skein across different developmental stages (**SI Fig. S2**). The granule cluster may dominate the cytoplasm and span across 80% of the apical-basal axis (**Movie S3**,**S4**). Fluorescence staining with eosin suggests that the ETC granules are composed of protein, but we have no information about the identity of the proteins.

### 2.3 Shape and size of epidermal threads

From transmission electron microscopy (TEM) and confocal microscopy, we found three levels of thread structures: (1) At the nanometer scale, TEM images show parallel filaments that are likely intermediate filaments, which is consistent with previous results (Blackstad, 1963; see **SI Fig. S3D**). (2) At the micrometer scale, epidermal threads trace regular right-handed helices. (3) At the sub-cellular scale, the helical thread is packed in a single layer in a switchback pattern against the inner plasma membrane surface (**Fig. 2C**; **Movie S4**). At one of its ends, it is interwoven among granules (**SI Fig. S2B**), which configuration may contribute to the scaffolding function once ETC contents are released (see below).

All epidermal threads examined were right-handed helices (*N* = 25 cells). To understand the helical geometry of threads, we randomly sampled helix sections with centerline lengths of 5 – 15 μm and found that the thread diameter (*ϕ*) varied between 0.2 – 1.0 μm (0.52 ± 0.18 μm; mean ± S.D.). The helical pitch angle (*θ*) varied between 47.6° – 81.8° (63.5° ± 5.6°) and was relatively consistent across the full thread diameter range for a given segment of thread. Similarly, the helical diameter (*D*) varied between 0.07 – 0.78 μm (0.35 ± 0.10 μm), with a slight reduction with increasing *ϕ* (**Fig. 2D**). The pitch angle allowed us to calculate how much the threads can increase in length if the helix is pulled taut. The extension can be characterized by an extension ratio *R*_*Ext*_ = 1 – *sinθ*, which averaged 10.1% over the range of pitch angles described above.

### 2.4 Epidermal threads versus slime threads

Due to the complex shape of threads, their long aspect ratios, and the difficulty of tracing threads among granules, we were not able to reconstruct the morphology of an entire thread using confocal microscopy. We were able, however, to measure the full length of threads we collected by scraping hagfish skin with a cover glass. Thread length *L*_*T*_ from these measurements was 2.2 ± 0.54 mm (mean ± S.D.; **SI Fig. S3A**,**B**). These isolated threads were generally straight and showed little evidence of the helical morphology seen in intact ETCs. Incorporating the helical pitch angle *θ* above, we can approximate the total length of the helical centerline as 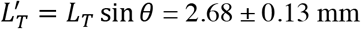, which is ∼53 times longer than the cell’s major axis (∼50 μm). Overall, the epidermal threads are ∼80 times shorter and ∼4 times thinner than slime threads, making them one of the largest intracellular fibers known (**Fig. 3B**). Some epidermal threads appeared to cleave into multiple sub-threads after being stretched, implying loose inter-filament binding (**Fig. 3C**; **SI Fig. S3C**).

**Figure 3.**
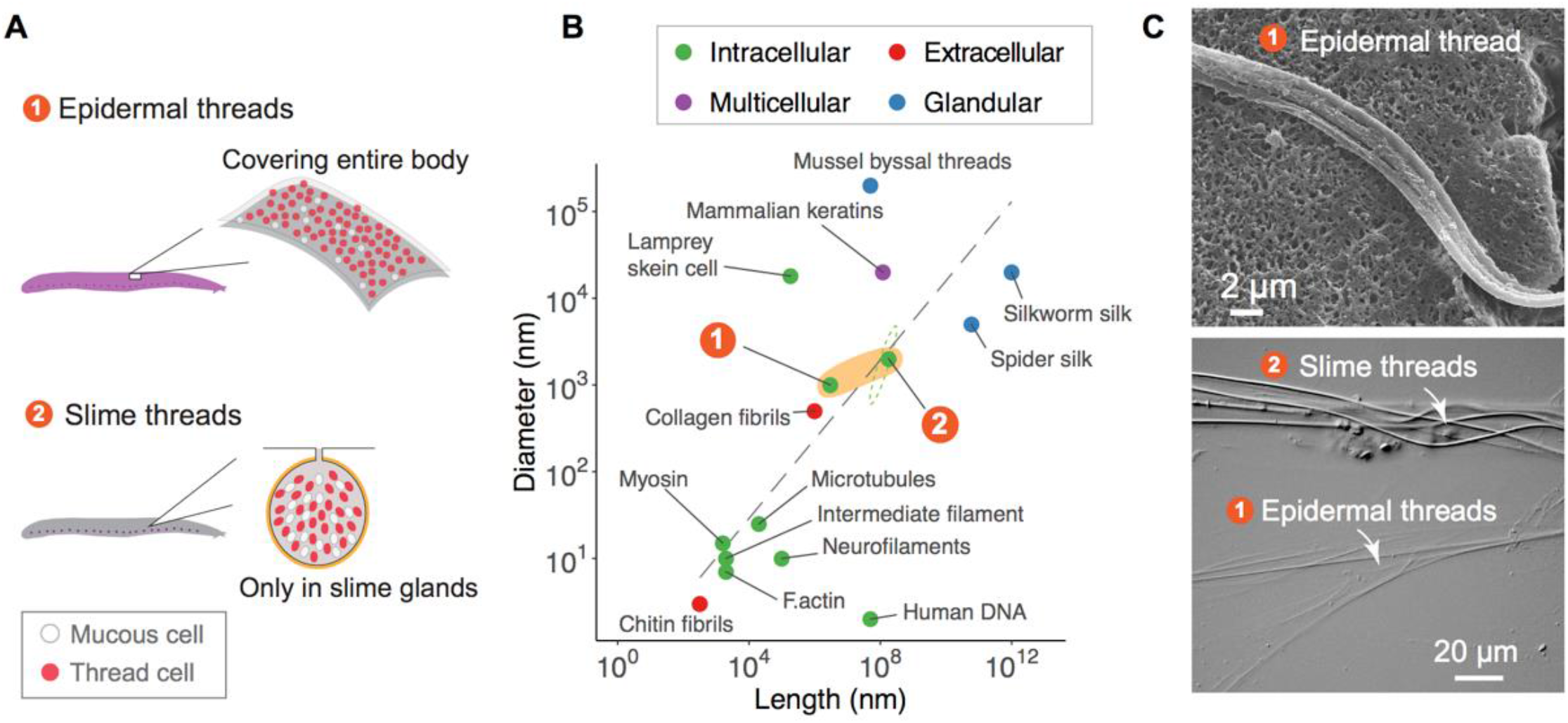
Comparison of distribution, size and shape between epidermal and slime threads. **(A)** A comparison of distribution between epidermal and slime threads. Note both types of thread cells are companioned by corresponding mucous cells in epiermis or slime glands. See **SI Fig. S1** for detailed analyses on epidermal cell abundance. **(B)** A size comparison of hagfish’s epidermal and slime threads (highlighted by orange shading) with other biofibers. The dashed ellipse near slime threads represents the full range of size variation in 19 hagfish species (length ∼5 cm to ∼22 cm; maximum diameter ∼0.7 μm to ∼3.9 μm). Trend line represents a linear regression model based on all data points excluding human DNA. Colors denote different fiber production mechanisms (see Zeng et al., 2021). **(C)** (Top) A section of an epidermal thread that has appeared to cleave into multiple sub-threads after being stretched, imaged with scanning electron microscopy (see also **SI Fig. S3C**). (Bottom) Two types of threads collected from the same hagfish (viewed with differential interference contrast microscopy), highlighting their difference in diameter.

Assuming threads are cylindrical and ETCs are ellipsoidal, the volume fraction occupied by threads within ETCs can be approximated as

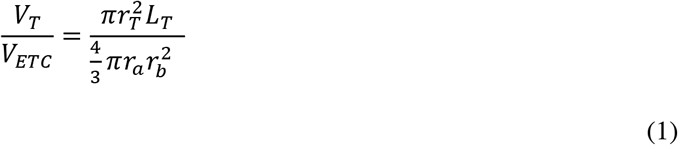

Using the ranges of thread radius (*r*_*T*_ = 0.5*ϕ*), thread length *L*_*T*_ and mean cell dimensions (major axis *r*_*a*_ ∼27 μm; minor axis *r*_*b*_ ∼ 23 μm; see **SI Fig. S2**), we found the epidermal threads only occupy 1.4% – 5.8% of the cytoplasmic space, which is much lower than the GTCs, where thread skeins may occupy > 95% of the cytoplasmic space (Downing et al., 1981; Zeng et al., 2021). To assess the thread storing capacity of the skin, we combined the stored thread length and area density of ETCs to calculate the area density of threads: *σ*_*T*_ = *σ*_*ETC*_*L*_*T*_, which yields a total of ∼96 cm threads per square millimeter of skin.

### 2.5 Damaged skin produces a fibrous slime

Dragging a sharp pin across a hagfish’s skin resulted in the formation and accumulation of a thick epidermal slime (translational speed ∼17 cm/s; mean vertical force 0.06 N, pressure ∼2 MPa, assuming a contact area of 0.03 mm^2^; **Fig. 4A,B**; **Movie S5**). Examining the path of the pin on the skin with scanning electron microscopy (SEM) revealed evidence that scraping caused rupture of ETCs and release of granules and threads. In relatively shallow wounds, where only the apical portions of ETCs were removed, the granule-thread complex was typically found anchored with the basal portion of threads to the inner surface of the cell’s plasma membrane (**Fig. 4C**).

**Figure 4.**
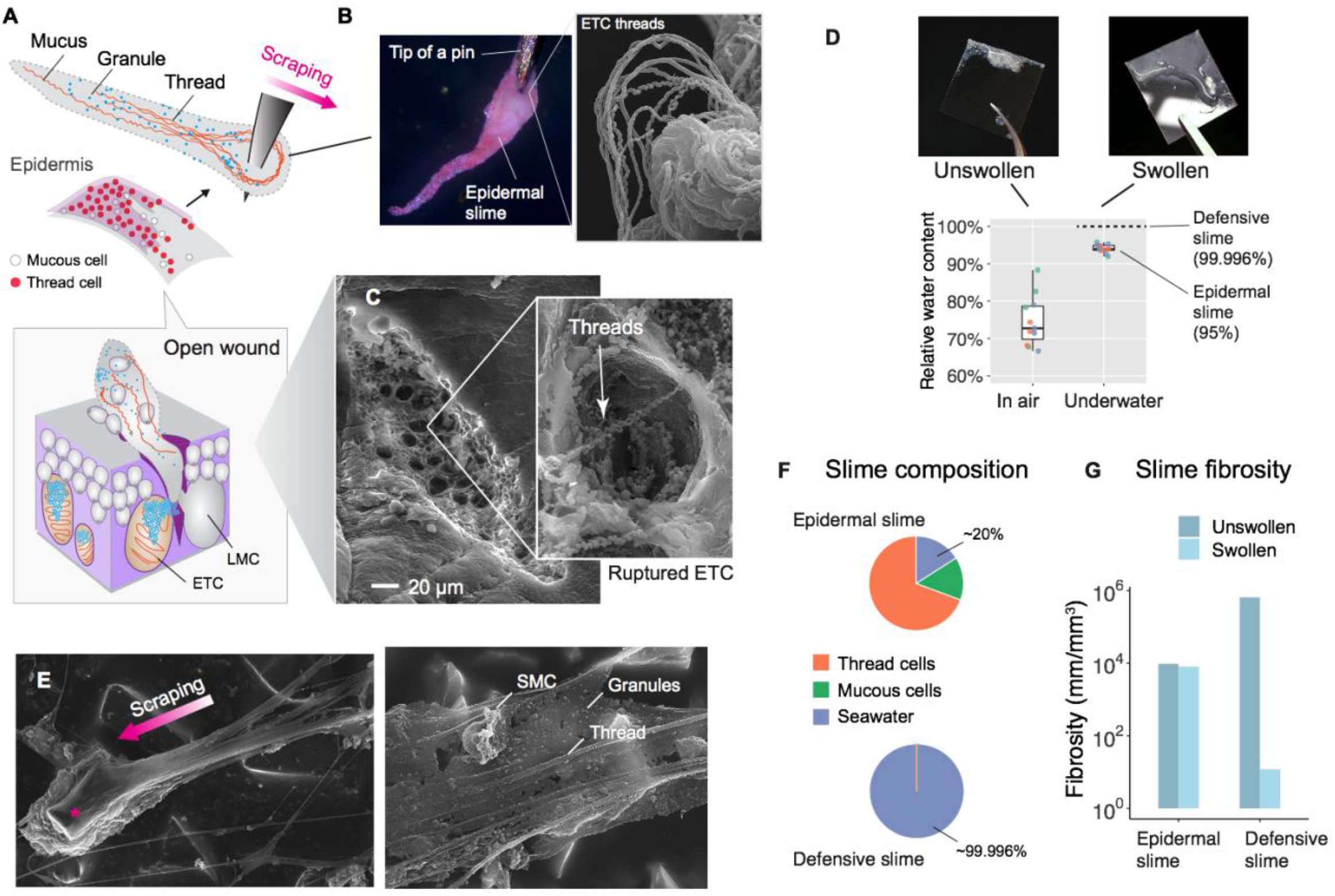
Formation and structure of epidermal slime produced by wounded skin. **(A)** Schematic of epidermal slime formation when epidermis is wounded, with threads and granules from ruptured ETCs mixing with mucus from ruptured LMCs. Bottom shows a schematic of the slime formation by mixing of cellular contents from an open wound on epidermis. **(B)** Epidermal slime on pin tip, stained with eosin to show threads. (Right) SEM image of epidermal slime on pin tip, with enlarged areas showing stretched and unstretched threads. **(C)** SEM images of a shallow abrasion wound, with insets showing damaged ETCs with partially released threads and granules. **(D)** The relative water content of epidermal slime collected by scraping a glass coverslip over blotted skin (unswollen) and underwater (swollen). Dots represent individual samples; colors represent different animals (*N* = 3 for each group; see **Methods**). See **SI Fig. S3-S5** for more information on epidermal threads and epidermal slime. **(E)** (Left) Epidermal slime collected on sandpaper. Note the slime accumulated at the leading edge of the sand grain and the elongated slime at the trailing edge. (Right) Thin film of epidermal slime collected by scraping with sandpaper, showing the scaffolding of mucus by threads, and the alignment of threads with the scraping direction. **(F)-(G)** A comparison of slime composition (in relative volumes) and fibrosity between epidermal and defensive slimes. Note the high water content and low fibrosity of defensive slime produced with turbulent mixing after active ejection. See **SI Table S2** for details.

Epidermal slime appeared as a white material that adhered to the scraping object, exhibiting properties distinct from the defensive slime (**Movie S5**,**S6**). Examination of the slime with light microscopy and SEM confirmed the presence of granules and threads, along with threads aligned with the scraping direction (**Fig. 4E**; **SI Fig. S4-S5**). Released granule-thread complexes were observed on the edge of coverslips used for scraping or on the skin surface after scraping, and often were seen with a single thread trailing from a granule cluster (**SI Fig. S3E-G**). Although we saw no direct evidence of LMC cell products, given their position in the same basal layer of the epidermis, it is likely that LMCs rupture under the same conditions that cause ETC rupture and contribute to the mucus components of epidermal slime.

### 2.6 Epidermal slime versus defensive slime

Scraping with the edge of a cover glass over 18 cm^2^ of skin that had been blotted dry led to about 2 – 10 mg (5.2 ± 2.4 mg; mean±S.D.) of slime adhered to the coverslip, which is equivalent to a productivity of ∼0.3 mg/cm^2^. The relative water content of epidermal slime sampled from skin immersed in seawater ranged from 92% – 96% (93.9% ± 1.2%; mean ± S.D.) and from 70% – 90% (74.7% ± 6.8%) for samples collected from dried skin (**Fig. 4D**).

Both epidermal slime and defensive slime are structurally heterogeneous, containing long threads and mucus. Here, we use the ratio between the total thread length and the total slime volume to characterize the level of ‘fibrosity’ of the two types of slime. The fibrosity index of epidermal slime was calculated as:

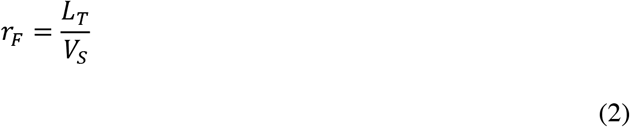

where *L*_*T*_ is the total length of thread and *V*_*S*_ is the volume of slime. Specifically, *L*_*T*_ was calculated as the product between the mean length of a single thread 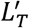 and the number of ETCs: *L*_*T*_ = *L*_*T*_*′N*_*ETC*_.

Considering an ideal situation without swelling with seawater, the volume of slime should equal to the total volume of ruptured epidermal cells. With a unit skin area *A* and the mean thickness of epidermis *D*_*epi*_ = 100 μm, we have *V*_*S(Unswollen)*_ = *AD*_*epi*_ and *N*_*ETC*_ = *σ*_*ETC*_*A*. Thus, Eqn. (2) can be expressed as:

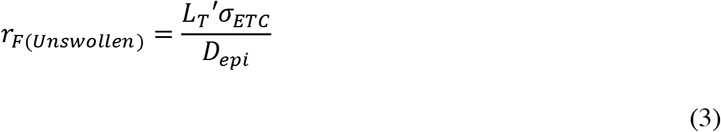

Incorporating single thread length (*L*_*T*_′ = 2.2 mm) and area density of ETC (*σ*_*ETC*_ = 434 mm^-2^), we found *r*_*F(Unswollen)*_≈ 9600 mm/mm^3^ for epidermal slime without swelling with seawater. Next, acknowledging that swollen slime has ∼19% more water than unswollen slime and assuming the density of unswollen slime is close to that of seawater, we derived *V*_*S*(*Swollen*)_ = 1.19*V*_*S(Unswollen)*_ and approximated *r*_*F*(*Swollen*)_ ≈ 8024 mm/mm^3^ for swollen epidermal slime, which is ∼686 times higher than that of the defensive slime (∼12 mm/mm^3^; based on Schorno et al., 2018; see **SI Appendix B**). Together, these results show that epidermal slime is less dilute and much more fibrous than defensive slime (**Fig. 4F,G**).

### 2.7 Epidermal threads are ancestral to slime threads

We examined the transcriptomes of skin and slime glands. Two types of thread proteins, α and γ, were previously identified (Koch et al. 1994; 1995) and threads produced from these genes were hypothesized to comprise the fibrous slime of hagfish. We characterized α and γ thread transcripts from replicate RNAseq datasets from skin and slime gland tissues of *E. goslinei*, a close relative of *E. stoutii*.

We identified a single, highly expressed α thread biopolymer transcript in the epidermis that likely comprises the epidermal threads (**Fig. 5**). We also uncovered a monophyletic diversity of highly expressed slime gland-specific α transcripts, suggesting that rampant gene duplication of GTC-specific α thread genes may underpin some of the exotic biophysical properties of hagfish slime. In addition, γ thread biopolymer transcripts in slime glands were more diverse than previously described and were only present in slime glands. The presence of well characterized, skin-specific *α* thread orthologs from both lamprey and teleosts indicates that a gene duplication of a skin-expressed *α* locus gave rise to a radiation of slime gland-specific *α* transcripts, while all *γ* biopolymer transcripts uniquely expressed in slime gland were secondarily derived.

**Figure 5.**
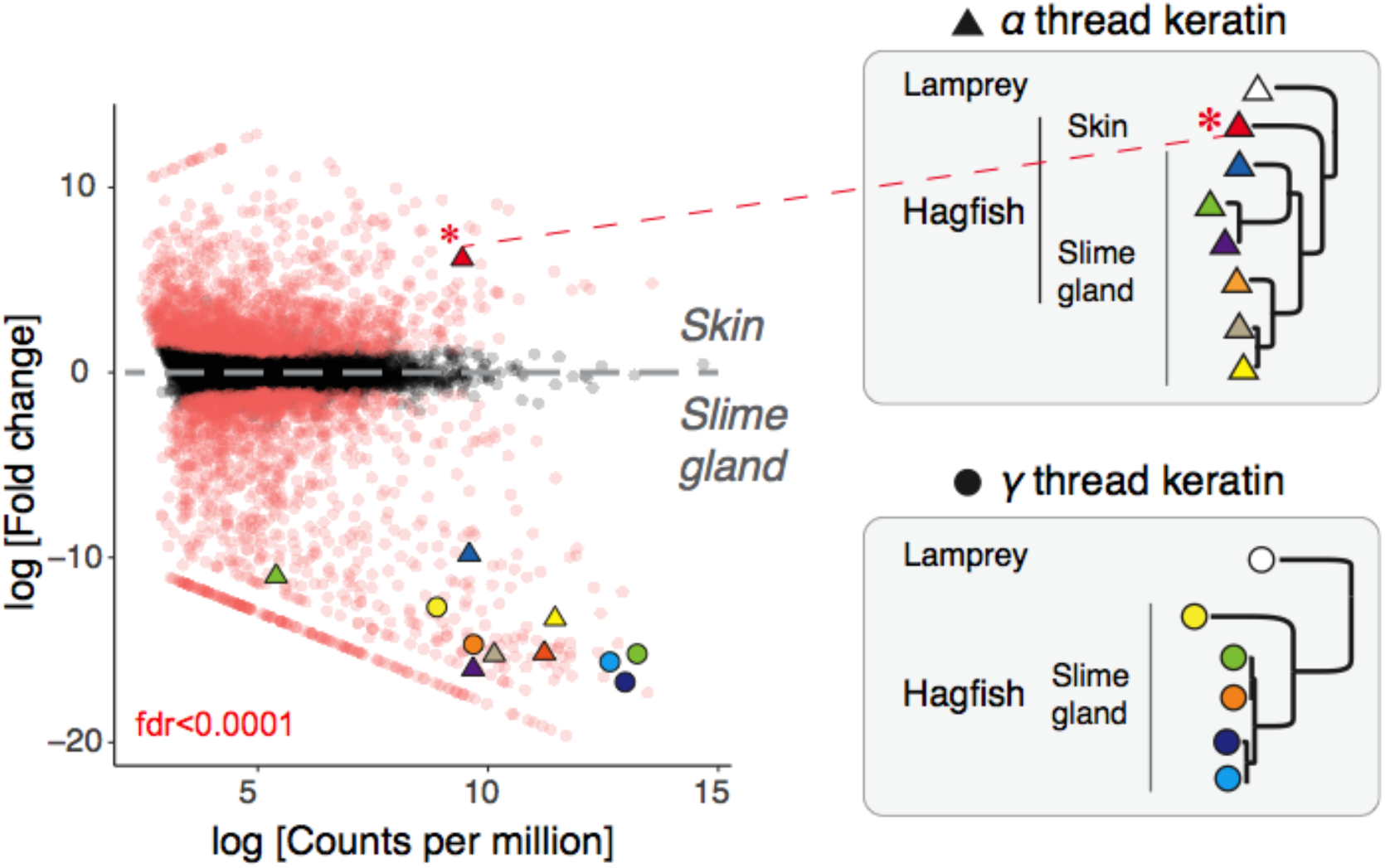
Molecular analyses suggest an epidermal origin of hagfish defensive slime. **(Left)** Differentially expressed (DE) transcripts (red) from skin vs. slime gland RNAseq read datasets (3× replicates each, single *E. goslinei* specimen; FDR < 0.001). A single *α* thread biopolymer gene is expressed in skin, while a diversity of both *α* and *γ* thread biopolymer genes are expressed in slime gland. **(Right)** Comparative phylogenomic analyses of *α* and *γ* thread gene trees (Maximum likelihood) identified slime gland- and hagfish-specific expansions of both *α* and *γ* intermediate filament genes. The presence of well characterized, skin-specific *α* thread orthologs from both lamprey and teleosts indicates that a gene duplication of a skin-expressed *α* locus gave rise to a radiation of slime gland-specific *α* transcripts. See **SI Fig. S6** for details.

## 3. Discussion

Our results demonstrated the epidermal threads are released through cell rupture during skin damage, together with mucus, forming a fibrous epidermal slime. With the ETC granules possibly serving anti-predator, antimicrobial or alarm functions (see **SI Appendix C**), the epidermal slime can be produced during interactions with predators and likely represent an incipient form of the defensive slime. Also, gene expression data provided support for an epidermal origin of slime glands. Below, we discuss the structure and function of epidermal threads and propose a simplified model to explain the origins of hagfish slime glands and defensive slime.

### 3.1 Structure and function of epidermal threads

Like slime threads, epidermal threads appeared to be mechanically robust, with no evidence of threads breaking even when they were sheared under a cover glass. The lack of helical twists in the elongated threads suggests that the threads are capable of plastic deformation, a property that has also been observed in slime threads and individual intermediate filaments (Fudge et al. 2003; Kreplak et al. 2005). Notably, the appearance of loose subfilament structure in some epidermal threads (**Fig. 3C; SI Fig. S3C**) has not been observed in slime threads. This suggests that epidermal threads may simply be a bundle of individual intermediate filaments. In contrast, intermediate filaments in slime threads undergo a phase transition in which filaments condense with their neighbors to form a single, solid thread (Winegard et al. 2014; Terakado et al., 1975; Downing et al., 1984).

The production of a macroscopic thread that is released after cell rupture suggests an evolutionary affinity between ETCs and GTCs and provides support for an epidermal origin of slime glands. If GTCs were derived from a primitive form of ETCs, selection for greater thread length and strength (and therefore diameter; **Fig. 3B**) were likely important for the transition from ETC to GTC. Selection for larger threads within the confined limits of the cytoplasm was also likely responsible for the evolution of a tightly packed thread skein and the loss of granules in GTCs (see Zeng et al., 2021). Hagfish GTCs and ETCs, along with lamprey skein cells (Land and Whitear, 1980), are the only epidermal cells capable of producing the largest intracellular fibers (**Fig. 3B**), and they likely share a single evolutionary origin.

Our results show that epidermal threads associate with mucus to form a fibrous epidermal slime, which may be the evolutionary precursor of defensive slime (see below). While the length of individual epidermal threads is small compared to slime threads, the large number of ETCs in the epidermis represents a significant reserve of thread length. For example, the total length of epidermal threads produced by ∼1.1% of the skin area of an adult hagfish (∼3.15 cm^2^) equals the total length of slime threads ejected from a single slime gland (∼2736 m) (see **SI Table S2**). These numbers demonstrate that a version of the most complex part of hagfish slime – the threads – was likely being produced in large quantities in the skin long before the slime glands appeared.

### 3.2 Mechanism of epidermal slime formation

Our mechanical abrasion experiments demonstrated the formation of a thick epidermal slime, which shared similar structural components with the defensive slime (**Fig. 6A**). Epidermal threads not only appeared to hold the slime together, but they were also readily caught on and adhered to the hard structures we used to damage hagfish skin (i.e., pin, coverslip and sandpaper; **Fig. 4**; **SI Fig. S5**). Under natural conditions, this property of the epidermal slime may allow it to adhere to a predator’s teeth after it has bitten a hagfish. Once bound to the predator’s teeth, the epidermal slime may deliver distasteful compounds to discourage further bites. It is also possible that the slime remains adhered to the hagfish’s skin after an attack (see **SI Fig. S4**), which would be consistent with an antimicrobial function of ETC granules, with compounds in the granules inhibiting bacterial growth at the wound site. Both the distasteful and antimicrobial hypotheses of epidermal slime function should be tested with further experiments.

**Figure 6.**
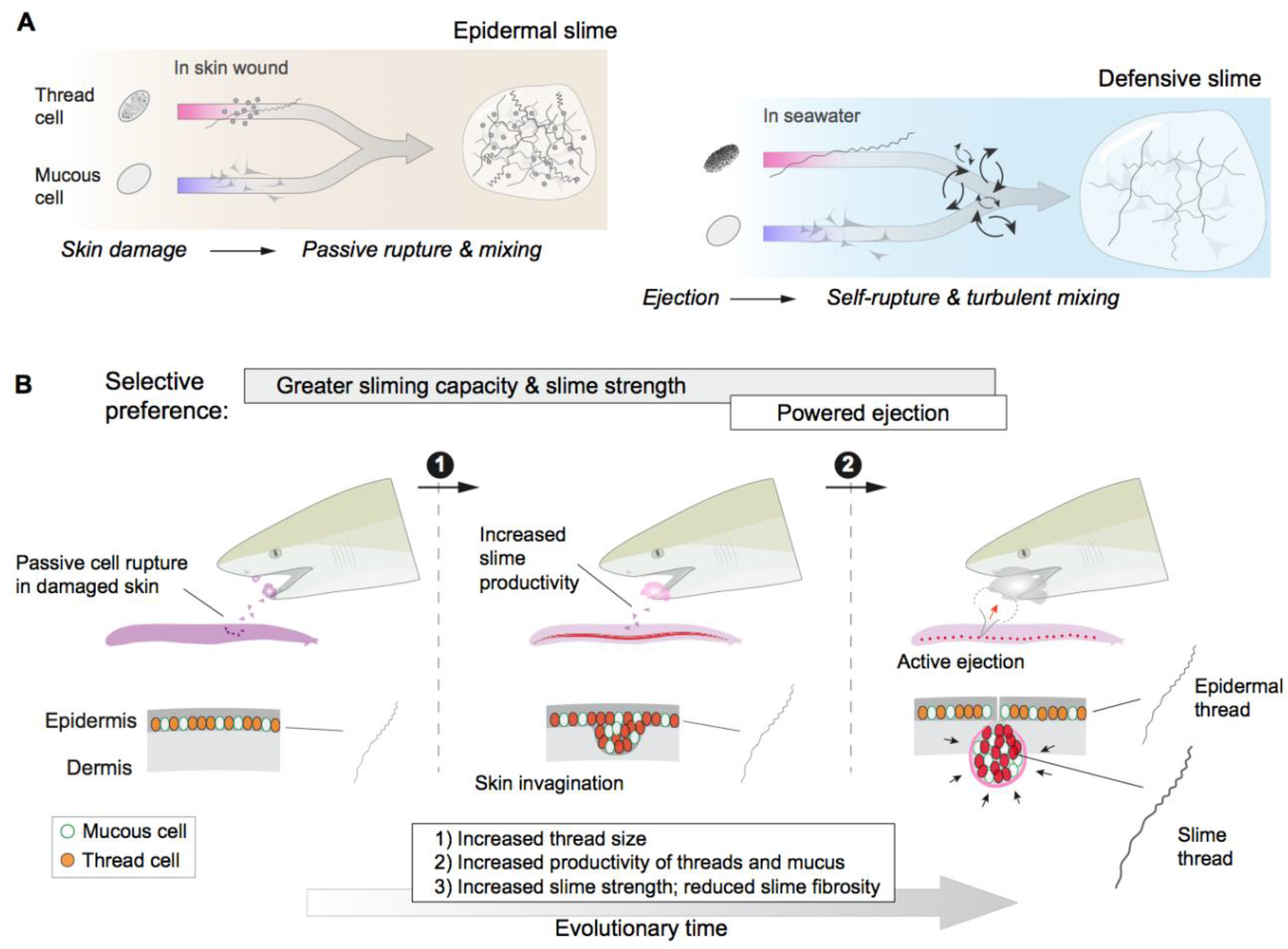
An epidermal origin of hagfish slime. **(A)** A comparison of slime formation mechanism between epidermal and defensive slimes, highlighting their similarity in basic structural components and differences in mixing mechanism. Note a transition from passive slime formation to active ejection, as well as a transition in slime composition. **(B)** Schematic of two critical transitions in the evolution of hagfish slime glands. Specifically, selection for greater slime capacity likely drove an increase in the concentration of thread cells and mucous cells in epidermis and later in slime glands, while selection for active ejection likely was responsible for the acquisition of gland musculature and an enlarged gland cavity with a narrow pore (see **Discussion**). Bottom row highlights the invagination of epidermis (middle) as a possible intermediate state between the ancestral form (left) and muscularized slime glands seen in modern hagfishes (right).

### 3.3 Thread proteins in skin and slime glands

Our transcriptomic analyses suggest that slime threads are evolved from epidermal threads, with duplication and diversification of skin-specific α genes and new expression of γ thread genes in slime glands (**Fig. 5**). If epidermal threads consist of bundled intermediate filaments made primarily of α thread proteins, this suggests that α proteins in the skin behave more like homopolymeric Type III intermediate filaments, whereas those in the slime glands are more similar to the heteropolymeric keratins (Type I/II). If true, this transition from a homopolymeric to a heteropolymeric intermediate filament may provide deep insights into how the keratin intermediate filaments may have arisen from homopolymeric ancestral proteins. It also reconciles the seemingly contradictory findings of Koch et al. (1994, 1995) and Schaffeld & Schultess (2006) who found similarities to Type I/II and Type III intermediate filaments, respectively.

To reduce heterozygosity, comparative transcriptome analyses were conducted using data from a single *E. goslinei* individual (Mincarone et al. 2021). In addition, we filtered our phylogenetic analyses of transcripts to include only approximately full-length sequences that had expression levels above TPM > 10 for replicate RNAseq datasets. Because of this, the diversity of thread transcripts identified from a single *E. goslinei* individual could correspond to prominently expressed loci, alleles, splice-products, and combinations therein. We screened publicly available coding sequence data from *E. burgeri* but did not detect sequences with homology to either α or γ. The lack of α and γ sequences in the *E. burgeri* genome may be a consequence of chromosome elimination, which has been shown to be prevalent in hagfishes (Nakai et al., 1995). While questions on the genetics of the α and γ thread diversity will become clear once more complete genomic resources for hagfish become available, the starkness of the expression differences between α and γ transcripts is notable. While we had no criteria that transcripts be differentially expressed between skin and slime gland for inclusion in our analysis, all α and γ transcripts that met the above criteria were significantly differentially expressed.

### 3.4 Implications for the origin of hagfish slime

The morphological, functional, and genetic evidence laid out above all point to an epidermal origin of hagfish slime glands. Below, we list some of the changes that had to occur to transform the epidermis into slime glands and we discuss the selective scenarios underlying those changes.

If one considers the origin of slime glands from a cellular perspective, there were several changes that had to occur, from the cellular composition and organization of the tissues, as well as changes to the cell themselves. Slime glands contain two main secretory cell types - GTCs and GMCs - and these most likely arose via modifications of ETCs and LMCs, respectively. For the transition from GTCs to ETCs, in addition to the increase in cell size (i.e., GTCs are ∼40 times larger than ETCs in volume) and thread packing due to selection for larger thread size (Zeng et al. 2021), some of the differences in thread properties may be related to the differences in thread protein composition that our transcriptome data point to, with slime glands expressing several α and γ transcripts and skin only expressing a single α transcript. While SMCs are the most common cell type in the epidermis, there is no corresponding cell type in the slime gland. The exclusion of SMC in slime glands probably has to do with its function of constitutively secreting mucus as a protective barrier at the outer surface of skin (Patzner et al., 1982). During the evolution of slime glands through possible invagination of the epidermis, cells specialized for slow release of mucus had little purpose and were likely excluded in favor of larger proportions of ETCs and LMCs (see below).

In addition to changes in cellular composition and the nature of the cells themselves, slime gland tissue differs markedly from epidermis, with the most obvious differences being their size, shape, and association with striated muscle. Slime glands are approximately ellipsoidal and typically 2-3 mm in diameter, whereas hagfish epidermis has a thickness of only about 100 µm. Thus, the production of slime glands from epidermis involved a local expansion of the epidermis, which was presumably driven by an initial selection for a greater capacity to produce mucus and threads (**Fig. 6B**). Expansion, and possibly invagination of the epidermis allowed for increased production and storage of secretory cells, but also created a new challenge of how to effectively deploy the secretory products. A selection for rapid release of a large number of thread and mucous cells (see also below) likely has fostered the acquisition of striated muscle fibers (i.e., the *musculus decussatus*) that surround the gland capsule. The muscle-powered ejection of gland cells allows hagfishes to produce large volumes of defensive slime at specific locations along their body in a few hundred milliseconds.

Our demonstration of the epidermal slime produced by damaged skin provides a possible starting point for addressing the initial selective scenarios underlying the origin of slime glands. We propose that hagfish epidermal slime arose as an immediate defense against predators, with distasteful granules released from ETCs acting to discourage further attacks. Under this scenario, epidermal threads may have arisen as a way of keeping the granules from dispersing too quickly after cell rupture. If this strategy was effective, selection may have favored an increased capacity to produce granules, threads, and mucus during attacks. At some point, the volume of released slime was enough for it to have other effects, most notably an ability to stick to the mouth and gills of fish predators due to the presence of the threads. We propose that it was this shift that ultimately led to the divergence of epidermis and slime glands, with threads in the former becoming specialized for binding granules and epidermal slime and threads in the latter specialized for clogging gills in association with mucus and seawater.

## 4. Materials and methods

### 4.1 Animal care and euthanasia

Wild-captured Pacific Hagfishes (*E. stoutii*) were housed in a 1000-liter tank of chilled artificial seawater (34%, 8°C) at Chapman University, CA, USA. Hagfish were anesthetized using clove oil (200 mg/L) (McCord et al. 2020). For euthanasia, hagfish were first anesthetized in 200 mg/L of clove oil and then transferred to a lethal dose of MS-222 (250 mg/L).

### 4.2 Abundance of epidermal thread cells

To quantify the abundance of ETCs and the other two epidermal cells, we sampled cell densities using fixed and stained samples of hagfish skin. First, with a series of transverse cross-sections, we sampled cell abundance along the skin circumference. One Pacific hagfish (body length ∼45cm) was fixed with 3% PBS-buffered paraformaldehyde, and then divided into 10 sections of equal length, exposing 9 transverse cross-sections. Of each cross-section, the anterior portion (∼1 cm thickness) was embedded in paraffin wax, sectioned (20 μm thick) and transferred to slides (**SI Fig. S1C**). The tissues were then stained with hematoxylin and eosin (H&E) following standard procedures (Bancroft and Gamble, 2008) and mounted with Permount Mounting Medium (Fisher SP15-100). Digital images were taken for the entire skin section using transmitted light microscopy (40× objective, Zeiss Axio Imager 2).

For each cross-section, the anteroposterior position (P_AP_) was defined as the relative distance from the snout (**SI Fig. S1C**). Next, we traced the profile of epidermis for one arbitrary side using ImageJ (Rueden et al., 2017). The dorsoventral position (P_DV_) was defined as the relative distance from the dorsalmost point (P_DV_=0; at the dorsal ridge). We then sampled sections of ∼1 mm long at each of dorsalmost, ventralmost and lateral positions. The dorsoventral position of each section was calculated as P_DV_=(P_b_-P_a_)/2, where P_a_ and P_b_ are dorsoventral positions of the two ends (**SI Fig. S1C**). Within each section, we manually recorded the number of cells (*N*_*cell*_) and calculated the linear density as *λ* = *N*_*cell*_/*L*_*section*_, where *L*_*section*_ = *P*_*b*_ − *P*_*a*_ is the section length. Analyses were performed using custom-written scripts in R (R Core Team, 2013).

Second, we sampled the area density (*σ*) of cells in 2 freshly euthanized hagfishes. From each hagfish, we collected skin samples (2×2 mm) from the lateral region at three anteroposterior positions (0.2, 0.5, and 0.8). Each skin sample was immediately fixed with 3% PBS-buffered paraformaldehyde (30 min), stained with eosin (∼2 min) and washed with 75% ethanol. The skin sample was then transferred to a large coverslip (24×50 mm) with the epidermis facing downward and covered by a smaller coverslip (24×40 mm). Images stacks were then taken with an inverted confocal microscope (Zeiss LSM 980).

### 4.3 Morphometrics of ETCs and contents

We took image stacks for ETCs on H&E stained slides using laser confocal microscopy (Zeiss LSM 980 with Airyscan). We sampled the size and area density of granules from cross-sectional confocal images of 17 ETCs. With each cross-section, we manually counted the number of granules and digitized the profile of the granule cluster. We then calculated the area density as *σ* = *N*/*A*, where *N* is number of granules and *A* is cross-sectional area of the cluster.

We further sampled granules from confocal image stacks taken in the axial direction to assess the variation of granule size. On each slice, we approximated each granule as an ellipse by fitting it with the ‘oval’ tool in ImageJ. We then summarized the size and density of granules with respect to the axial position (as represented by z-direction) using custom-written R scripts.

### 4.4 Size and shape of epidermal threads

The helical geometry of threads was sampled from confocal image stacks using ImageJ. We chose helix sections that revolved about an approximately straight central axis for at least 3 consecutive helical loops. We also checked the thread appearance between stacks to make sure it was approximately aligned with the image plane. We placed paired landmarks on the peaks and valleys on each side of the thread section (**SI Fig. S2C**). Later, with custom-written R scripts, we calculated the centerline of each helix as *p*_*c*_ = <*p*_*i*_ + *p*_*j*_ >, where *p*_*i*_ and *p*_*j*_ denote points on each bilateral side of the thread and angle brackets denote average. The mean direction of increase was represented by a vector 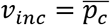. The thread diameter (*ϕ*) was calculated as *ϕ* = |*p*_*i*_ − *p*_*j*_| for each pair of landmarks and the mean diameter was calculated for each helix. The pitch angle (*θ*) was calculated for each half loop as the angle between the centerline and the normal direction of the mean direction of increase. Correspondingly, the helical radius (*r*) was calculated as 2*r* = *p*/tan*θ*, where *p* is the helical pitch angle (**Fig. 2D**).

### 4.5 Epidermis wounds

We examined the products of epidermal abrasion caused by frictional contact and laceration caused by sharp surfaces. To simulate the frictional contact with epidermis and collect the products, we scraped the epidermis of anesthetized hagfishes using a glass coverslip (18×18 mm). In each trial, we oriented the coverslip at a ∼45° contact angle to the hagfish skin and scraped along the lateral side for a linear distance of < 5 cm. Next, the coverslip was carefully placed onto a glass slide (**SI Fig. S3E**). The samples were then observed with an upright compound microscope using transmitted light and DIC optics (Zeiss Axio Imager 2) and images were captured with a digital camera (Axiocam 506; 2752 × 2208 pixels). For free threads, we took individual images with 20× or 40× objective lenses and later stitched them using Adobe Photoshop.

To observe wounded epidermis, we introduced shallow wounds with a scalpel on euthanized hagfishes. We then excised a 2×2 mm skin sample and placed each on a large coverslip (24×50 mm) with the epidermis facing down. The samples were fixed with 4% PBS-buffered paraformaldehyde (∼20 min), stained with eosin (∼5 min) and washed with 75% ethanol. To minimize disruption, the samples were maintained on the coverslip throughout the staining process. We washed the samples by slightly tilting the coverslip and dropping 75% ethanol from the higher end, with paper towel collecting the liquid at the bottom. We then took images of the samples using confocal microscopy (Zeiss LSM 980 with Airyscan).

### 4.6 Phylogenetic and comparative transcriptome analyses

Transcriptome assemblies were constructed using Trinity (Grabherr et al., 2011) using RNAseq datasets from three replicates of skin and slime gland tissues for *E. goslinei*. Resulting assemblies were filtered using cd-hit and a -c 0.98 parameter setting. Reduced transcriptome assemblies were then translated to protein sequences using Transdecoder (Grabherr et al., 2011). Concurrently, reads from the replicate RNAseq datasets were mapped onto the assemblies using Salmon (Patro et al., 2017) and differential gene expression analyses were conducted using the Fisher’s Exact test implemented in EdgeR (Robinson et al., 2010) with *P*-value cutoff of 0.05.

Database searching and phylogenetic analyses were conducted using the following approach. First, a BLAST (Altschul et al., 1990) database was prepared that included protein models from the genomes of *Petromyzon marinus, Callorhinchus milli*, and *Danio rerio*, and the translated protein models derived from the *E. goslinei* transcriptome assembly. BLAST (Altschul et al., 1990) was conducted using α and γ thread sequences (Koch et al., 1995) as queries in separate analyses using a low stringency e value of 0.0001 while retaining up to 30 sequences per species. The resulting sequences were aligned using MAFTT (Katoh and Standley, 2013) and the –auto setting and the first round of phylogenetic analyses were conducted under the best fit model in IQ-TREE (Nguyen et al., 2015), which in both cases was LG+I+G+F. The resulting topologies, after rooting with a distant intermediate filament outgroup, contained many additional, more distantly related intermediate filament proteins in addition to the α and γ thread clades. Next, in separate phylogenetic analyses of α and γ sequences, approximately full-length sequences that had an expression of transcripts per million (TPM) > 10 were retained as were any non-hagfish sequences that may have been present. Finally, in separate procedures α and γ sequences were realigned and analyzed phylogenetically under the best fit model (LG+I+G+F) resulting in the gene trees shown in **SI Fig. S6**. Bioinformatic and statistical code is available at https://github.com/plachetzki/ETC_GTC. Raw RNAseq data are available under BioProject PRJNA497829.

## Acknowledgements

We thank Andrew Lowe for logistical help and thank Richard Wassersug for comments. This study was supported by NSF grants IOS-1755397 to D.F. and IOS-1755337 to D.P.

## Author contributions

Y. Z., K. N., H. C., and D. F. devised experiments. Y. Z., K. N., H. C. and K. G. collected and analyzed morphological and experimental data. D.P. and M. C. conducted the molecular and phylogenetic analyses. Y.Z., D.F., and D.P. all contributed to writing the manuscript.

## Declaration of interests

The authors declare no competing interests.

## Notes

https://github.com/plachetzki/ETC_GTC

### Competing Interest Statement

The authors have declared no competing interest.

