## Supplementary figures for "Epidermal threads reveal the origin of hagfish slime"

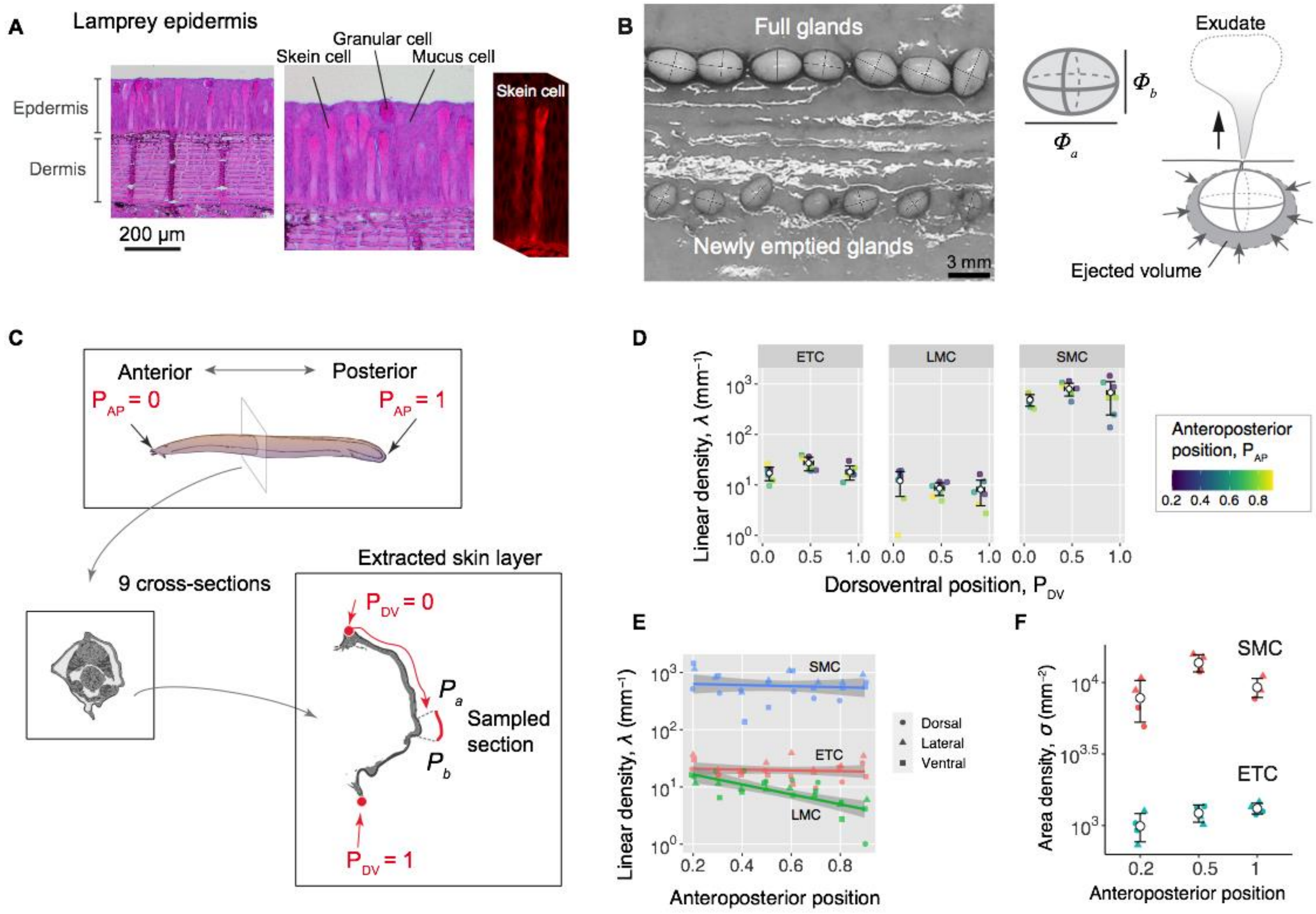


**Figure S1. Morphology of thread cells and slime gland.**
**(A)** Cross-section of lamprey (*Petromyzon marinus*) epidermis based on H&E-stained cross-sectional slides. Right shows a lamprey skein cell imaged with confocal laser scanning microscopy. The large fiber in lamprey skein cell ranges 80–120 µm and 30–60 µm in length and width, respectively (see also Lane and Whitear, 1980).
**(B)** Morphometrics of hagfish slime gland. **(Left)** Full glands and newly emptied glands on the same hagfish, showing difference in size and similarity in shape (see Schorno et al., 2018). We found that full and newly emptied glands share a similar aspect ratio of ~1.44 (see **SI Appendix A**). **(Right)** Configuration of major and minor axes for a slime gland. The volume of ejected exudate per gland can be approximated as the volume difference between full and newly emptied glands (see **SI Appendix B**).
**(C)-(F)** Abundance of hagfish epidermal cells. **(C)** Schematic showing anteroposterior position ($P_{AP}$), with 0 representing the snout and 1 the tip of the tail. (Bottom left) Schematic showing dorsoventral positions ($P_{DV}$) in a hagfish in cross-section (see Methods). **(D)** Linear density ($\lambda$) of each cell type for dorsal, lateral and ventral regions along the hagfish ($N$ = 1 hagfish). Values are means±S.D. Dots represent data for each cross-section, with color representing $P_{AP}$. The linear density of ETCs was relatively consistent as a function of circumferential position, with the mean value ranging between 17 – 27 $mm^{-1}$. The other two cell types, SMCs and LMCs, had similarly uniform distributions. The SMCs exhibited the highest density (126 – 434 $mm^{-1}$), and the LMCs had the lowest among the three (2 – 6 $mm^{-1}$). **(E)** Among all three types of epidermal cells, only the LMCs exhibited a change in density with respect to the anteroposterior position. Based on linear regression models, we found no significant effect of anteroposterior position on the densities of ETCs and SMCs ($P > 0.1$). However, in LMCs, we found a significant effect, with slope = -1.47±0.30, $P < 0.001$. **(F)** Area density ($\sigma$) of SMCs and ETCs sampled from laser confocal images taken in *en face* view from two hagfishes at three different anteroposterior positions ($N$ = 2 samples per location). Values are means±S.D. Colored dots represent data for each sampled area.

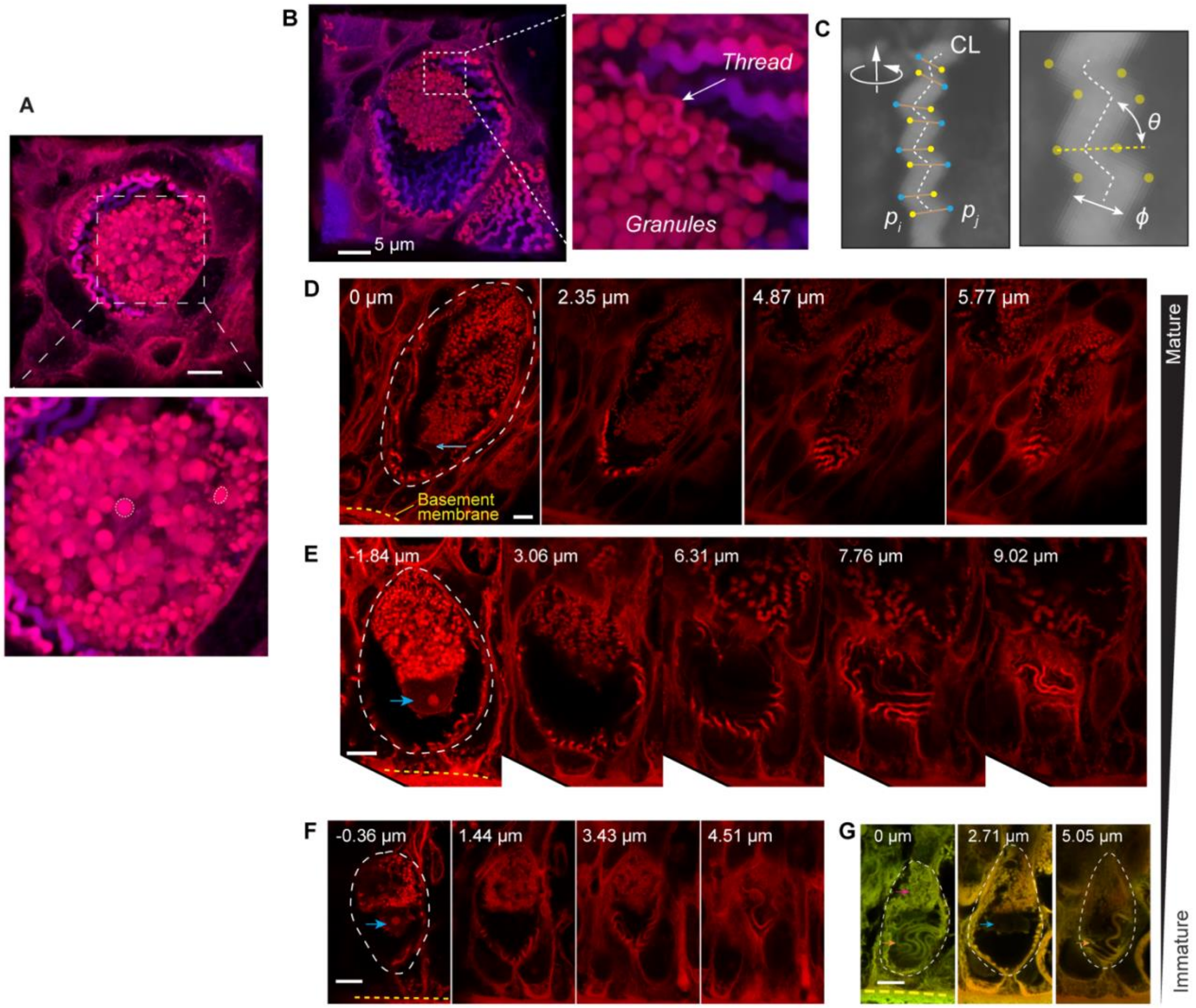


**Figure S2. Morphology of epidermal thread cell (ETC) from confocal laser scanning microscopy.**
**(A)** Confocal microscopic image showing the granule cluster in axial view (from exterior toward the animal). Enlarged area shows variations of granule size and shape.
**(B)** Oblique cross-section of an ETC, showing the relative positions of the granule cluster and threads. Enlarged area shows a region where the thread is intimately associated with the granule cluster.
**(C)** The peaks and valleys of the projected thread sections were used as landmarks for morphometric analysis. Blue dots, peaks; yellow dots, valleys; white dashed line, centerline. $\phi$, thread diameter; $\theta$, helical pitch angle; $D$, helical diameter.
**(D)-(G)** Developmental sequence of ETCs represented by cells of different sizes, with smaller cells at the bottom. Each cell is shown with images stacks at different z-distance, as annotated on the top. Note the variations in thread shape. Scale bars, 5 μm. All images were acquired from H&E stained slides using confocal microscopy (Zeiss LSM980). The mature ETCs are 54.1 ± 2.8 μm in length (in apical-basal direction) and 22.8 ± 0.6 μm in width (means±S.D.; $N$ = 5 cells).

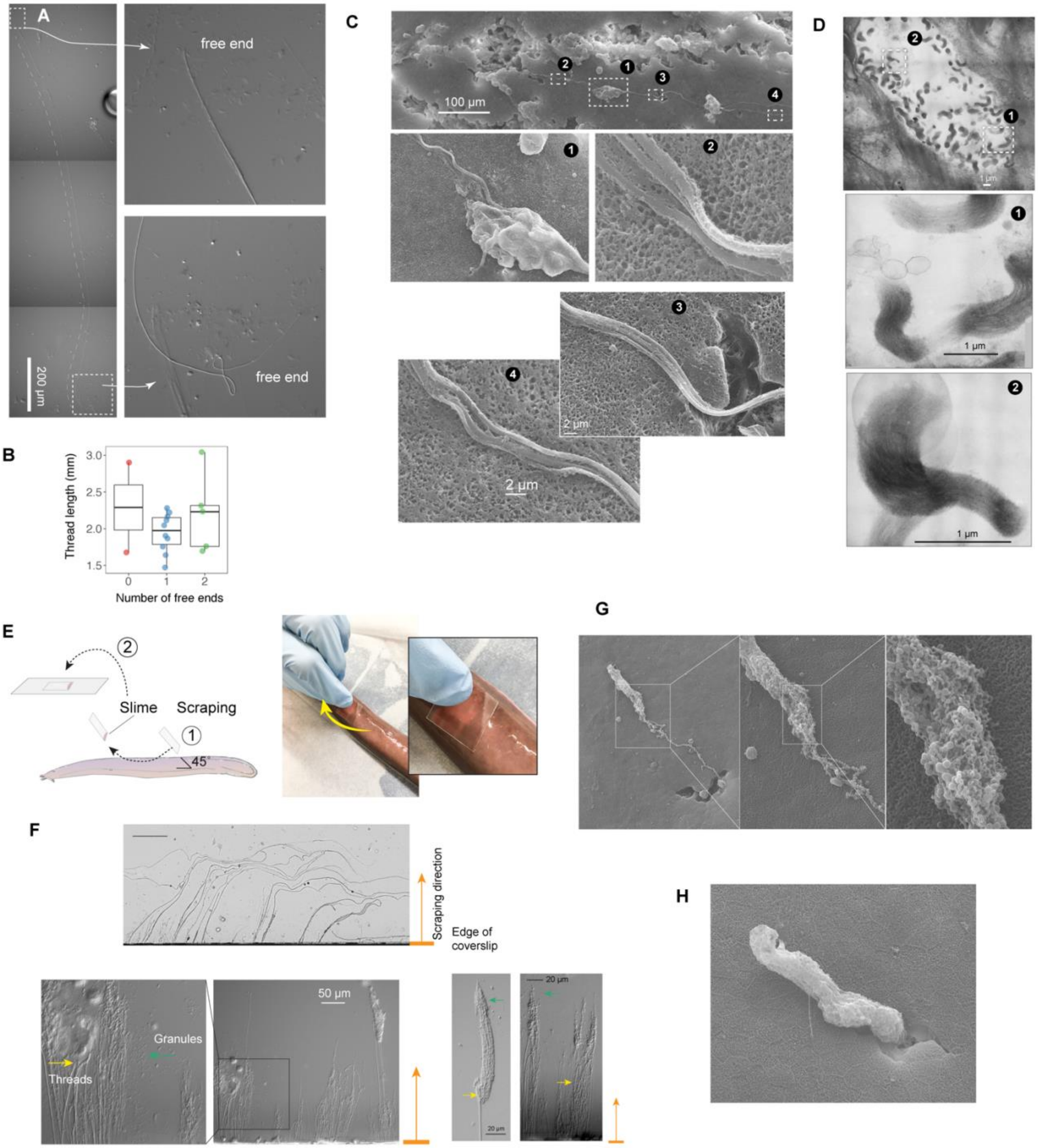


**Figure S3. Morphology and release of epidermal threads.**
Observing epidermal slime samples from scraping experiments under light microscopy, we found the epidermal threads lacked obvious helicity as seen in threads within cells, suggesting possible plastic deformation of threads during cell rupture. **(A)** shows a thread with two free ends. **(B)** Boxplots of thread length ($L_T$) measurements based on threads with no free end ($N$ = 2), one free end ($N$ = 10), and two free ends ($N$ = 5). Values are length from individual threads. For threads with one free end, $L_T$ = 1.95±0.27 mm (mean±S.D.); for threads with two free ends, $L_T$ = 2.2±0.54 mm, with this mean value used in scaling models. **(C)** SEM images showing details of a single ETC thread on the epidermal surface abraded with sandpaper, showing subfilament structure.
**(D)** Transmission electron microscopy (TEM) of an ETC in cross-section, showing sub-micron structures of threads as parallelly aligned filament bundles, which is consistent with the thread structure observed in the Atlantic hagfish (*Myxine glutinosa*) (Blackstad, 1963).

**(E)** Schematic of sampling epidermal mucus using a coverslip scraped along the skin of an anesthetized hagfish. The coverslip was only gently pressed against the hagfish skin, as shown on the right.
**(F)** ETC threads captured by the edge of coverslips, as viewed under light microscopy.
After scraping hagfish skin with the edge of a coverslip, ruptured ETCs and free granules and threads were evident near the coverslip edge. Enlarged area shows threads and granules. The bottom row shows ruptured ETCs with unraveled threads.
**(G)** A granule cluster found on the epidermal surface, with a helical thread still attached. **(H)** Partially released ETC granule cluster found on the epidermal surface.

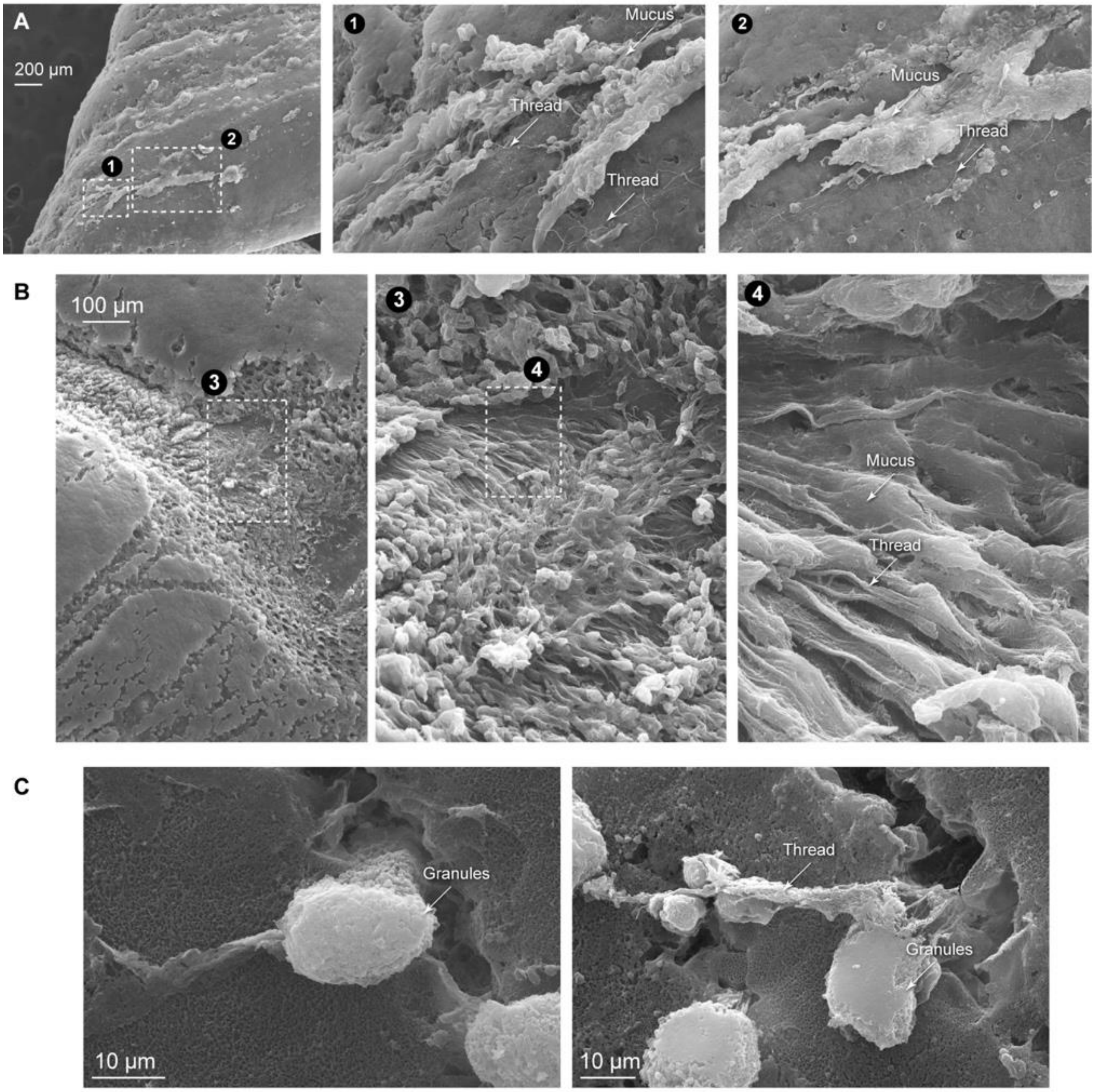


**Figure S4. Formation of epidermal slime.** Newly wounded hagfish skin was fixed, dehydrated and observed using scanning electron microscopy (SEM). **(A)-(B)** Formation of epidermal slime on abraded skin, with details showing the slime as a mixture of mucus, ETC threads and SMCs. **(C)** The release of ETC granules and threads from laceration wounds made with a scalpel.

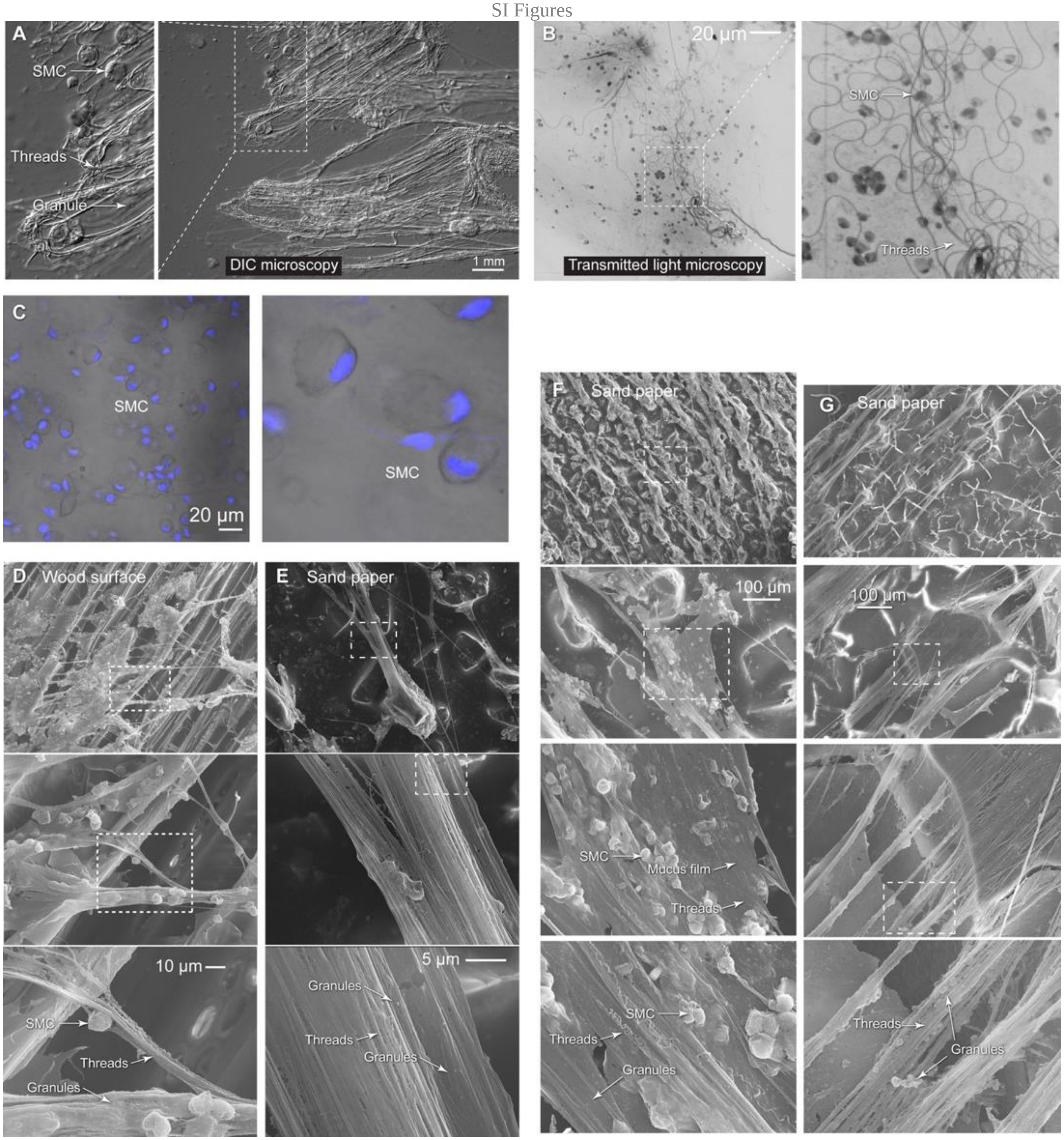


**Figure S5. Structure of epidermal slime.**
**(A)** Epidermal slime observed using differential interference contrast (DIC) microscopy, showing ETC threads (yellow arrow), granules (green arrow) and detached SMCs (purple arrow).
**(B)** Epidermal slime collected after scraping hagfish skin with a pin and observed using bright-field microscopy, showing individual ETC threads.
**(C)** SMCs appeared mostly intact in epidermal slime, although may have been ruptured. Here we show SMCs with nuclei highlighted using DAPI.
**(D)-(G)** After scraping hagfish skin with rough surfaces (sand paper and wood), we observed epidermal slime adhered to these surfaces using SEM. The epidermal slime appeared as thin strings or sheets adhered to the protruding surfaces, with the rest trailing or crossing over gaps. From top to bottom, images show details of the epidermal slime with progressively increasing magnification. Dashed boxes denote enlarged areas.

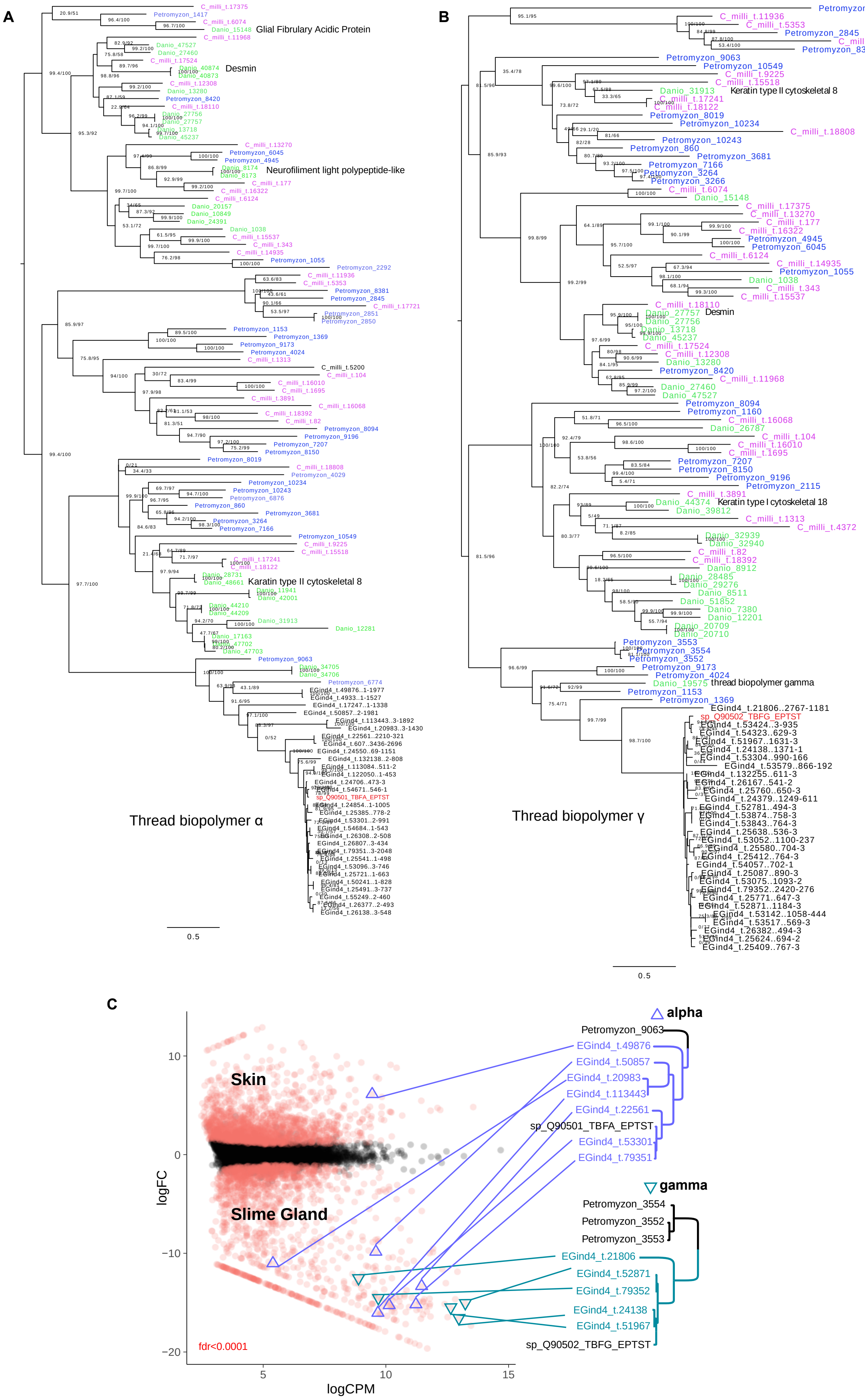
A
Glial Fibrulary Acidic Protein
Desmin
Neurofiliment light polypeptide-like
Karatin type II cytoskeletal 8
Thread biopolymer α
0.5
B
Keratin type II cytoskeletal 8
Desmin
Keratin type I cytoskeletal 18
thread biopolymer gamma
Thread biopolymer γ
0.5
C
alpha
Petromyzon_9063
EGind4_t.49876
EGind4_t.50857
EGind4_t.20983
EGind4_t.113443
EGind4_t.22561
sp_Q90501_TBFA_EPTST
EGind4_t.53301
EGind4_t.79351
gamma
Petromyzon_3554
Petromyzon_3552
Petromyzon_3553
EGind4_t.21806
EGind4_t.52871
EGind4_t.79352
EGind4_t.24138
EGind4_t.51967
sp_Q90502_TBFG_EPTST
Skin
Slime Gland
fdr<0.0001
logFC
logCPM
10
0
-10
-20
5
10
15

**Figure S6. Molecular Analysis support an epidermal origin of hagfish defensive slime.**
**(A)-(B)** Comparative phylogenomic analyses of α and γ thread gene trees (Maximum likelihood, best fit; L+G+I+F; Nguyen et al., 2015) identifies slime gland- and hagfish-specific expansions of both α and γ intermediate filament genes. The presence of well characterized, skin-specific alpha thread orthologs from both lamprey and teleosts (Schaffeld & Schultess, 2006) indicates that a gene duplication of a skin-expressed alpha locus gave rise a radiation of slime gland-specific α transcripts. All γ biopolymer transcripts recovered in this analysis were expressed uniquely in slime gland GTCs. Selected sequences from *Danio rerio* are annotated.
**(C)** DE transcripts (red) from skin vs. slime gland RNAseq read datasets (3× replicates each, single *E. goslinei* specimen; FDR<0.001 Robinson et al., 2010). A single α thread biopolymer gene is expressed in skin ETCs, while a diversity of both α and γ thread biopolymer genes are expressed in slime gland GTCs. All assembled transcripts show in **(A)** and **(B)**. Only transcripts that had expression greater than 10 TPM shown in **(C)**.
