## Supplementary text for "Epidermal threads reveal the origin of hagfish slime"

### Supplementary Information

#### Movies

**S1.** Z-stack image sequences of eosin-stained hagfish epidermis from confocal laser scanning microscopy with transmitted light, taken in *en face* view. Note a dense layer of epidermal thread cells (ETCs) and large mucous cells (LMCs) at the basal layer of epidermis. Each ETC is evident with a cluster of granules highlighted in red, while LMCs appear as voids.

**S2.** Z-stack image sequences of hagfish epidermis from confocal laser scanning microscopy, taken in *en face* view. Note the outermost epidermis is covered by a layer of small mucus cells (SMCs), while ETCs are found at the basal layer.

**S3.** Z-stack image sequences of hagfish ETCs based on confocal laser scanning microscopy. Images were taken from eosin-stained epidermis, and the ETC granules are the brightest feature.

**S4.** Three-dimensional reconstructions of ETCs based on confocal laser scanning microscopy, showing granule cluster and helical-shaped threads packed along the plasma membrane.

**S5.** Experimentally induced formation of epidermal slime, demonstrated by scraping wet and blot-dried hagfish skin with a sharp pin head. Scraping with a blunt pinhead did not lead to slime formation.

**S6.** Different from condensed, adhesive hagfish epidermal slime, the defensive slime is highly diluted and non-sticky.

### Appendix

#### A. Abundance of GTCs in slime glands and exudate

**Slime gland shape is conserved after exudate ejection**. We found there was no significant difference in the aspect ratio (AR; i.e., the ratio of major axis to minor axis) between full glands and newly emptied glands. Using image data from a previous study (Schorno et al., 2018), we found that newly emptied glands are ~30% smaller in both major and minor axes than full glands (full glands: major axis $\Phi_a$ = 3.51±0.25 mm, minor axis $\Phi_b$ = 2.45±0.15 mm, *N* = 11; newly emptied glands: $\Phi_a$ = 2.47±0.25 mm; $\Phi_b$ = 1.71±0.17 mm, *N* = 10; means±S.D.; **SI Fig. S1B**). The gland AR was 1.43±0.09 for full glands and 1.45±0.13 for newly emptied glands. The slime glands were then modeled as ellipsoids with a mean AR = 1.44 (see below).

**Total number of slime glands**. In this study, we dissected one Pacific Hagfish (*Eptatretus stoutii*; body length, 48 cm), and counted the glands from under the skin. We found a total of 81 glands on the left side and 82 glands on the right side. A total of 163 glands was then used in assessing GTC abundance and productivity.

**Number of GTCs in full glands.** We assumed that GTCs are distributed evenly within the glands, which allowed us to estimate the total based on cross-sectional images. For the full glands, we approximated the total number of GTCs ($N_{GTC}$) based on the area density ($\sigma_{GTC}$) and gland volume ($V_G$) as $N_{GTC} = \sigma_{GTC}^{1.5} V_G$, where $V_G = \frac{4}{3}\pi r_a r_b^2$ and $r_a = 0.5\Phi_a$ and $r_b = 0.5\Phi_b$ are major and minor radius, respectively. Applying mean values of $\sigma_{GTC}$ and gland dimensions from literature (**Table S1**), we found there are ~19300 GTCs in each full gland. With 163 full glands, the total number of GTCs is ~$3.15 \times 10^6$, which is ~26% of the total number of ETCs in the epidermis (~$1.23 \times 10^7$, with the hagfish simplified as a cylinder of 45 cm in length and 20 cm in diameter).

**Number of GTCs ejected to seawater.** Similarly, with the mean values of $\sigma_{GTC}$ and gland dimensions from newly emptied glands, we found ~4100 GTCs remained in emptied gland after ejection. Subtracting the number of remaining GTCs from the total in full glands, we approximated that ~15200 GTCs are ejected per gland (**Table S1**).

**Volume of ejected exudate.** The volume of ejected exudate can be calculated as the volumetric difference between full gland and newly emptied gland as: $V_{[ejected]} = V_{G[full]} - V_{G[emptied]}$. We found 9.4 mm$^3$ exudate was ejected by each gland.

**Volume of ejected mucin.** The area proportions ($\delta$) of gland mucous cells (GMCs) in full and newly emptied glands were also available (Schorno et al., 2018). By assuming even distribution of GMCs within slime glands, we then calculated the volume of mucin as: $V_{GMC} = \delta_{GMC} V_G$ for both full and newly emptied glands. The ejected volume of GMCs was then calculated as $V_{GMC[ejected]} = V_{GMC[Full]} - V_{GMC[Emptied]}$.

#### B. Fibrosity of hagfish defensive slime

In this study, the 'fibrosity index' represents the ratio of total thread length in a soft material to the total (e.g., mucus and seawater for defensive slime; **SI Fig. S1B**). To approximate the fibrosity index of defensive slime, we used (1) the total length of GTC threads ejected by one slime gland and (2) an approximation of the total volume of mucus and seawater mixed with these threads, as derived below.

With ejected exudate volume $V_{[ejected]}$ = $9.37 \times 10^{-3}$ ml per gland and exudate density $\rho$ ~ 1 g/ml (a conservative estimate based on Fudge et al., 2005), we estimated the weight of ejected exudate to be $W = \rho V_{[ejected]}$ ~$9.37 \times 10^{-3}$ g. Also, the combined w/v concentration of thread and mucin is 0.004% in fully-deployed slime (Fudge et al., 2005), the volume of seawater mixed with the exudate from a single gland can then be estimated as: $V_{SW} = V_{[ejected]}/0.004\%$ = $9.37 \times 10^{-3}$ / 0.004% = 234.25 ml. Next, the total volume of liquid in fully-deployed slime is then $V_S = V_{SW} + V_{[ejected]}$ = 234254.2 mm$^3$ ≈ $2.34 \times 10^5$ mm$^3$. Lastly, the fibrosity of defensive slime was then calculated as:

$$r_F = \frac{L_T N_{GTC}}{V_S} \quad \text{(S1)}$$

We found the fibrosity is ~$6.5 \times 10^5$ mm/mm$^3$ for unmixed exudate and ~12 mm/mm$^3$ for fully-deployed defensive slime (**Table S2**), which shows that the exudate is diluted $5.5 \times 10^4$ times and that the fully-deployed defensive slime is ~800 times less fibrous than epidermal slime.

**Table S1. Morphological parameters of slime glands of Pacific Hagfish.** Values correspond with one full exhaustion of a single gland. †, mean values of morphological variables of full and newly emptied slime glands from literature (Schorno et al., 2018). ‡, single GTC thread length 18 cm was assumed (see Zeng et al., 2021).

| | Full gland | Newly emptied gland | Ejected exudate |
|---|---|---|---|
| Minor axis, $\Phi_a$ (mm) † | 2.62 | 1.77 | / |
| Major axis, $\Phi_b$ (mm) † | 3.77 | 2.55 | / |
| Volume (mm$^3$) | 13.6 | 4.2 | 9.4 |
| GTC area density, σ (mm$^{-2}$) | 126.5 | 98.8 | / |
| Total GTC number, $N_{GTC}$ | 19300 | 4100 | 15200 |
| Total thread length (m) ‡ | 3474 | 738 | 2736 |
| GMC area proportion, $\delta$ (%) † | 56.6 | 83.5 | / |
| GMC volume (mm$^3$) | 7.7 | 3.5 | 4.2 |

**Table S2. Comparison of fibrosity between defensive slime and epidermal slime.** Values for defensive slime correspond with one full exhaustion of a single gland. Values for epidermal slime correspond with 1 mm$^2$ of skin surface, with volume approximated based on the thickness of epidermis and experimentally measured relative water content (see main text).

| | Ejected exudate per gland | Fully-deployed slime per gland | Epidermal slime per 1 mm$^2$ skin |
|---|---|---|---|
| Total thread length (mm) | 2736000 | 2736000 | 955 |
| Volume (mm$^3$) | 9.4 | 234000 | 0.12 |
| Fibrosity index (mm/mm$^3$) | 651429 | 12 | 8024 |

### C. Possible functions of ETC granules

Within cross-sections of granule clusters, granule density was 1.05 ± 0.50 $\mu m^{-2}$ (mean±S.D.). Although generally round, the granules were not spherical. The aspect ratio (i.e., the ratio of major to minor axis) was 1.5 ± 0.6 (mean±S.D.; $N$ = 1462 granules).

While the thread is the most conspicuous part of an ETC when viewed with conventional histology and microscopy (hence their name), our measurements based on high-resolution confocal microscopy show that the granules take up far more volume in the cell than the thread. The production of granules in large numbers is typical of secretory cells (Bowen, 1929) and further suggests a secretory function served by ETCs (Blackstad, 1963; Spitzer and Koch, 1998). The lack of any obvious secretory mechanism for ETCs and the results of our skin wounding experiments lead us to the conclusion that granules are primarily released when the epidermis is damaged and ETCs are ruptured. In light of these results, we consider three possible functions for the granules: (1) They may contain distasteful compounds that help deter predators when hagfishes are bitten. A similar defensive function has been suggested for the epidermal granule cells of lampreys (Pfeiffer and Pletcher, 1964) and this would be a reasonable adaptation for hagfishes, whose scavenging lifestyle involves frequent bites from predators (Zinzten et al. 2011; Boggett et al. 2017). (2) The granules may contain anti-microbial compounds that help prevent infection after the skin is damaged (e.g., 'myxinidin'; Subramanian et al., 2009). This too would be sensible for an animal that is frequently bitten. (3) The granules may contain an alarm pheromone that alerts other hagfishes to the presence of an attacking predator, a mechanism that has been widely reported in lampreys and fishes (Bals and Wagner, 2012; Imre et al., 2014; Pandey et al., 2021). The identity of the proteins that make up ETC granules is unknown, but future work to identify and characterize these proteins will undoubtedly shed additional light on their function.
